# A Novel Technique to Characterize *Klebsiella pneumoniae* Populations Indicates that Mono-Colonization is Associated with Risk of Infection

**DOI:** 10.1101/2025.11.05.686704

**Authors:** Lavinia V. Unverdorben, Sophia Mason, Weisheng Wu, Moraima Noda, Stuart Castaneda, Jay Vornhagen, Evan S. Snitkin, Krishna Rao, Michael A. Bachman

## Abstract

*Klebsiella pneumoniae* and related species are a common cause of healthcare-associated infections. The gut is a major *Klebsiella* reservoir and gut colonization is a risk factor for developing an extraintestinal *Klebsiella* infection. Patients can be colonized by multiple *Klebsiella* strains or even species in the gut simultaneously, and there is high concordance between the gut colonizing- and infection causing-strains. The detection and characterization of colonizing strains is critical for a better understanding of the progression to infection and for developing interventions for colonized patients. However, the association between mixed or mono-colonization and subsequent infection is unknown. In this study, we developed an amplicon-based sequencing method called *wzi*-Seq that enables the detection and quantification of *Klebsiella* strains from complex samples and mixtures using the conserved capsule gene *wzi* as a molecular barcode. This method is highly accurate and precise with a sensitivity of 93% and specificity of 99.8% in mixtures containing as many as 58 unique *wzi* types. The assay was validated analytically and applied to an established case and control cohort. We determined that 63.2% (108/171) patients were mono-colonized with a single *Klebsiella* strain while 36.8% (63/171) had mixed colonization with multiple *Klebsiella* strains.

Controlling for patient variables in multivariate analysis, we determined that mono-colonization was significantly (p = 0.034) associated with infection. Characterization of *Klebsiella* colonizing populations could improve the accuracy of assessing infection risk and enable targeted interventions to prevent these healthcare-associated infections.

**Importance:** *Klebsiella* gut colonization is a major risk factor the development of extraintestinal *Klebsiella* infections in hospitalized settings. However, it is unknown if patients are colonized by one or multiple strains of *Klebsiella* and how the population structure of *Klebsiella* in the gut impacts infection risk. Here we describe the development of a novel technique, *wzi*-Seq, to detect and quantify multiple *Klebsiella* strains from complex samples using standard techniques and rapid DNA sequencing. We applied *wzi*-Seq to rectal swabs from a well-characterized patient cohort and found that *Klebsiella* colonization with a single strain was more prevalent than colonization with multiple strains. Furthermore, mono-colonized patients had a significantly higher risk of developing a *Klebsiella* infection than mixed colonized patients. This work validates a new tool to study *Klebsiella* populations and reveals that population structure in the gut influences the risk of healthcare associated infections.

## Introduction

The Gram-negative bacterium *Klebsiella pneumoniae* (*Kp*) has become a pathogen of major concern due to increasing virulence, infections, and antibiotic resistance^1,2^. The rise in *Kp* antibiotic resistance is especially concerning as *Kp* has become the third highest cause of deaths attributable or associated to antimicrobial resistance globally^3^. *Kp* is part of a species complex (KpSC) of five closely related species and their subspecies: *K. pneumoniae*, *K. variicola, K. quasipneumoniae*, *K. quasivariicola*, and *K. africana*^4^. These species can colonize mucosal surfaces in the body such as the nasopharynx and gut and can cause pneumonia, urinary tract infections, and bacteremia^5,6^.

One of the primary *Kp* reservoirs is the gut with asymptomatic gut colonization rates typically ranging between 6-23% ^7–10^. *Kp* gut colonization is a major risk factor for the development of an extraintestinal *Kp* infection^11,12^. In approximately 80% of patients there is concordance between the gut colonizing and infection causing strain^8,13^. Colonization data is based typically on a few cultured rectal isolates per patient, preventing a clear understanding of colonizing *Kp* dynamics in patients. Even with this limited sampling, we occasionally observed that patients were colonized with more than one strain simultaneously^13^. The true incidence of mixed colonization in patients, and its impact on infection risk is a major gap in knowledge. For example, mixed colonization could represent a low-risk state in which commensalism between strains, and the larger community prevents progression to disease, or an extremely dysbiotic state where niche exclusion is compromised and multiple pathogenic strains of *Kp* invade and increase the risk of infection.

Molecular typing schemes to identify and track *Kp* strains, including multilocus sequence typing (MLST), pulsed-field gel electrophoresis (PFGE), whole genome sequencing (WGS), and capsule typing are highly standardized and reproducible but are often time and labor intensive^14–18^. In recent years, strain typing based on the conserved *wzi* gene in the *Kp* capsule operon has become an alternative method for distinguishing *Kp* strains^16^. Previous work has validated *wzi* primers for the conserved motifs that flank the highly variable 447 bp region of the *wzi* gene (Fig. 1)^16^. Sanger sequencing of the *Kp wzi* gene using these primers has enabled the curation of the BigsDB database (https://bigsdb.pasteur.fr/klebsiella/klebsiella.html) containing 741 unique *wzi* types^16,19^. Strain identification via *wzi* typing has been shown to have higher discriminatory power than traditional K typing and similar discriminatory power as MLST typing^16^. This technique has become a fast and reliable alternative method for the differentiation of *Kp* strains.

**Figure 1:**
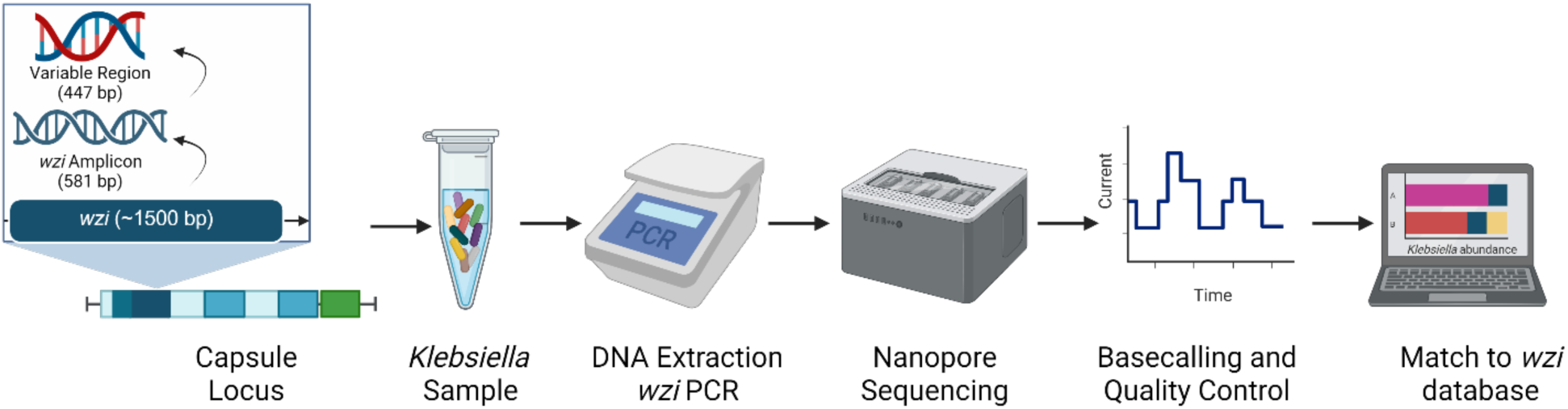
Overview of the *wzi*-Seq method A 581 bp *wzi* amplicon generated using previously published *wzi* primers, and the 447 bp variable region of the *wzi* amplicon are used to distinguish different *wzi* types. The *wzi*-Seq method starts with a *Klebsiella* sample with known or unknown *wzi* types which is then subject to DNA extraction and *wzi* PCR. Nanopore sequencing of the *wzi* amplicon is performed, followed by base calling and quality control of reads to ensure read accuracy. All reads that pass quality control and filtering are then aligned against a database of 741 known *wzi* type reference sequences to calculate the abundance of each *Klebsiella wzi* type in the sample.

To better characterize *Kp* colonization dynamics and the impact on patient infection risk we adapted *wzi* sequencing primers to develop a high throughput and accurate amplicon-based method called *wzi*-Seq^16^. The goal of this study was to develop and validate the *wzi*-Seq method, use it to assess the incidence of mono-vs mixed colonization in patients, and evaluate the impact of colonization status on patient infection risk. We demonstrated that *wzi*-Seq is a highly accurate and precise method with a sensitivity of 93% and specificity of 99.8% for detecting strains in complex mixtures containing up to 58 *wzi* types. We applied *wzi*-Seq to patient rectal swabs from a previous case-control study and demonstrated that mono-*Kp* colonization was more prevalent than mixed *Kp* colonization and associated with an increased risk of developing a subsequent extraintestinal *Kp* infection^13,20^. This work describes a novel method of using the *wzi* gene as a molecular barcode and provides insight into *Kp* colonization dynamics that has the potential to improve our understanding of *Kp* colonization dynamics and patient infection risk.

## Results

### wzi-Seq Assay Design

To design an amplicon-based sequencing method for the detection and quantification of multiple *Kp* strains from complex samples, we developed a nanopore-based long-read amplicon sequencing technique targeting the conserved *wzi* gene (Fig. 1). Sequencing reads were mapped to a reference database of 741 unique *wzi* alleles, and two quality filtering pipelines were used, which differed in allowing no mismatches (error0) or an error rate of up to 0.5% of the alignment length (error0.005). Whereas the error0 pipeline was more stringent for the presence of a strain, the error0.005 pipeline was expected to produce more reads for quantification of abundance. After quality filtering, the remaining reads for each *wzi* type were used to calculate the total abundance of each *wzi* type in a sample.

### Accuracy and Precision

To validate the *wzi*-Seq approach, contrived samples were created using increasingly complex mixtures of strains with known *wzi* types. *Kp* strains with unique *wzi* types were mixed at known ratios to create 1, 5,12, 30 or 58 *wzi* type sample mixtures. Sequencing of a single *Kp* strain KPPR1 (*wzi* type 2), yielded 13,367 and 6,181 reads from the error0.005 and error0 pipelines, respectively. Of these reads, the total correct mapped reads were 13,338 and 6,179 for the error0.005 and error0 pipelines, with a read accuracy greater than 99.7% for both pipelines (Suppl. Table 1). Although both pipelines had reads that mapped to at least two additional *wzi* types that were not the expected *wzi* type, these reads were less than 0.3% of the total for either pipeline (Suppl. Table 1).

To further test the accuracy and precision of the *wzi*-Seq method, a 5 *wzi* type mixture was created using *Kp* strains with *wzi* types 2, 57, 19, 522, 322. The *wzi* types were mixed using intended ratios of 50%, 30%, 15%, 4%, and 1% of each strain based on optical density (OD) measurements. An average of 9,932 and 4,322 reads were recovered from the error0.005 and error0 pipelines, respectively with a read accuracy greater than 99% for both pipelines (Suppl. Table 2). For the error0.005 pipeline, the mean *wzi* type abundance was within 1% of the expected abundance and there was high precision between replicates (standard deviation <1% for each *wzi* type; Table 1). Although the error0 pipeline had higher total read accuracy, the pipeline was slightly less precise (standard deviation: 0.1-1.1%) compared to the error0.005 pipeline (standard deviation: 0.1-0.6%) (Table 1). When comparing the experimental abundance values between the two pipelines there were no significant differences in the abundance values for each *wzi* type (Fig. 2A).

**Figure 2:**
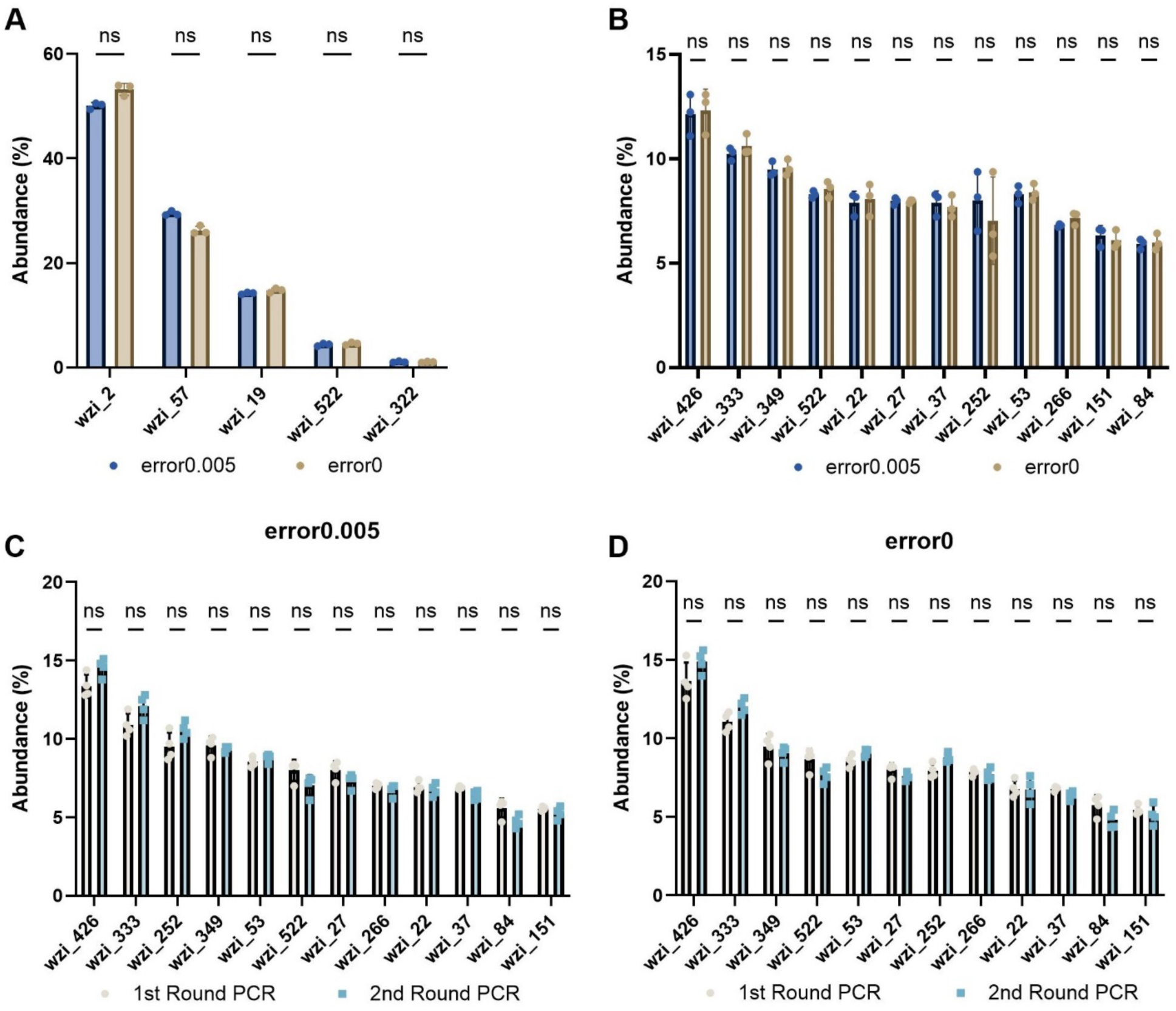
Abundance values are equivalent across pipelines and rounds of PCR (A) *wzi*-Seq was performed on a mixture of 5 *wzi* types wzi_2, wzi_57, wzi_19, wzi_522, and wzi_322 with expected relative abundances of 50, 30, 15, 5, and 1%, respectively. The mean abundance of each *wzi* type was determined from three replicates using the error0 and error0.005 pipelines. The abundance of each *wzi* type was compared between the two pipelines and there were no significant differences in the abundance for each *wzi* type as determined by the error0.005 or error0 pipelines. (B) *wzi*-Seq was performed on a mixture of 12 unique *wzi* types and the mean abundance of each *wzi* type was quantified by either the error0 or error0.005 pipeline. (C) A 12 *wzi* type mixture was subject to either one or two rounds of *wzi* PCR, with each round consisting of 40 cycles. The 12 *wzi* type mixtures subject to one or two rounds of PCR were analyzed using both the error0 (C) and error0.005 (D) pipelines to calculate the abundance of each *wzi* type. For (A-D), a multiple Wilcoxon paired t-test was used with an n≥3.

**Table 1:** Summary table of expected and experimental abundance values for 5 *wzi* type samples based on pipeline and rounds of PCR

| Pipeline |  | error0.005 |  | error0 |  | error0.005 |  |  |  |
| --- | --- | --- | --- | --- | --- | --- | --- | --- | --- |
| PCR |  | 1 Round |  | 1 Round |  | 1 Round |  | 2 Rounds |  |
| *wzi* Type | Expected Abundance (%) | Mean Abundance (%) | Standard Deviation | Mean Abundance (%) | Standard Deviation | Mean Abundance (%) | Standard Deviation | Mean Abundance (%) | Standard Deviation |
| 2 | 50 | 50.1 | 0.6 | 53.2 | 1.1 | 48.8 | 0.36 | 45.9 | 0.20 |
| 57 | 30 | 29.5 | 0.4 | 26.3 | 0.8 | 32.1 | 0.41 | 35.2 | 0.17 |
| 19 | 15 | 14.2 | 0.2 | 14.8 | 0.4 | 13.7 | 0.16 | 11.5 | 0.04 |
| 522 | 4 | 4.4 | 0.3 | 4.6 | 0.2 | 4.0 | 0.04 | 4.0 | 0.05 |
| 322 | 1 | 1.1 | 0.1 | 1.1 | 0.1 | 1.1 | 0.14 | 1.9 | 0.05 |

Next, we tested more complex mixtures containing between 12-58 unique *wzi* types. Sequencing a 12 *wzi* type mixture in triplicate yielded 96,296 and 47,064 correctly mapped reads for the error0.005 and error0 pipelines, respectively (Suppl. Table 2), for a read accuracy greater than 99%. There was no significant difference in the recovered abundance values for each *wzi* type between the two pipelines (Fig. 2B). Similarly to the 5 *wzi* type mixture, the precision between replicates was high with a standard deviation of less than 2% for the abundance of any of the 12 *wzi* types (Suppl. Table 3). Although more than 12 different *wzi* types were recovered by both pipelines, these erroneous reads made up less than 0.5% of all total reads and there was no significant difference in the abundance of each *wzi* type between the two pipelines (Fig. 2B, Suppl. Table 2). Additionally, the *wzi*-Seq method was tested in duplicate on 30 and 58 *wzi* type mixtures (Suppl. Table 4). The read accuracy was greater than 99% for the error0 pipeline, and 95.5% and 97.0% for the 30 and 58 *wzi* type mixtures, respectively, with the error0.005 pipeline (Suppl. Table 4). Together these data highlight that *wzi*-Seq can accurately differentiate *wzi* types in complex mixtures containing up to 58 unique *wzi* types.

To enable analysis of low-biomass samples in which *Kp* may be present in low abundance, the accuracy and precision of performing a second round of PCR was tested on 5 and 12 *wzi* type mixtures. There were no significant differences in the abundance values after one or two rounds of PCR in the 12 *wzi* type mixture regardless of whether the error0.005 or error0 pipeline was used (Fig. 2C, D). No differences between the number of rounds PCR and abundance levels were also observed for the 5 *wzi* type sample mixture (Table 1). Together these data highlight the high accuracy and precision of the *wzi*-Seq method and that this accuracy and precision is maintained with increasingly complex mixtures and low biomass samples.

### Sensitivity and Specificity of Strain Detection

Next, we assessed the true positive and false positive rates of strain detection in contrived mixtures. The *wzi* abundance values of 23 different contrived samples with mixtures ranging in complexity from 5-58 *wzi* types were used as the input to generate a receiver operating characteristic (ROC) curve. All detected *wzi* types that were expected in each sample were considered true positive values and all other *wzi* types were considered false positives. For the error0.005 and error0 pipelines, the area under the ROC was 0.9925 and 0.9986, indicating very high accuracy (Fig. 3A, B). The ROC curves were used to determine cutoffs for both pipelines that would optimize sensitivity and specificity of strain detection at the readfrac level before converting to relative abundance. When using a readfrac cutoff of 0.00275 on error0.005 samples, the predicted sensitivity and specificity of the assay was 93.0% and 99.8%, respectively (Fig. 3C). For the error0 pipeline, a readfrac cutoff of 0.00172 predicted an assay sensitivity and specificity of 94.2% and 100%, respectively (Fig. 3C). These cutoffs were then tested on four contrived mixtures containing between 5-30 different *wzi* types that were not included in the original ROC analysis. Applying these cutoffs eliminated 98.9% of false positives in error0.005 samples and 100% of false positives in error0 samples with 100% of true positives detected in all samples (data not shown) and were applied in subsequent analyses.

**Figure 3:**
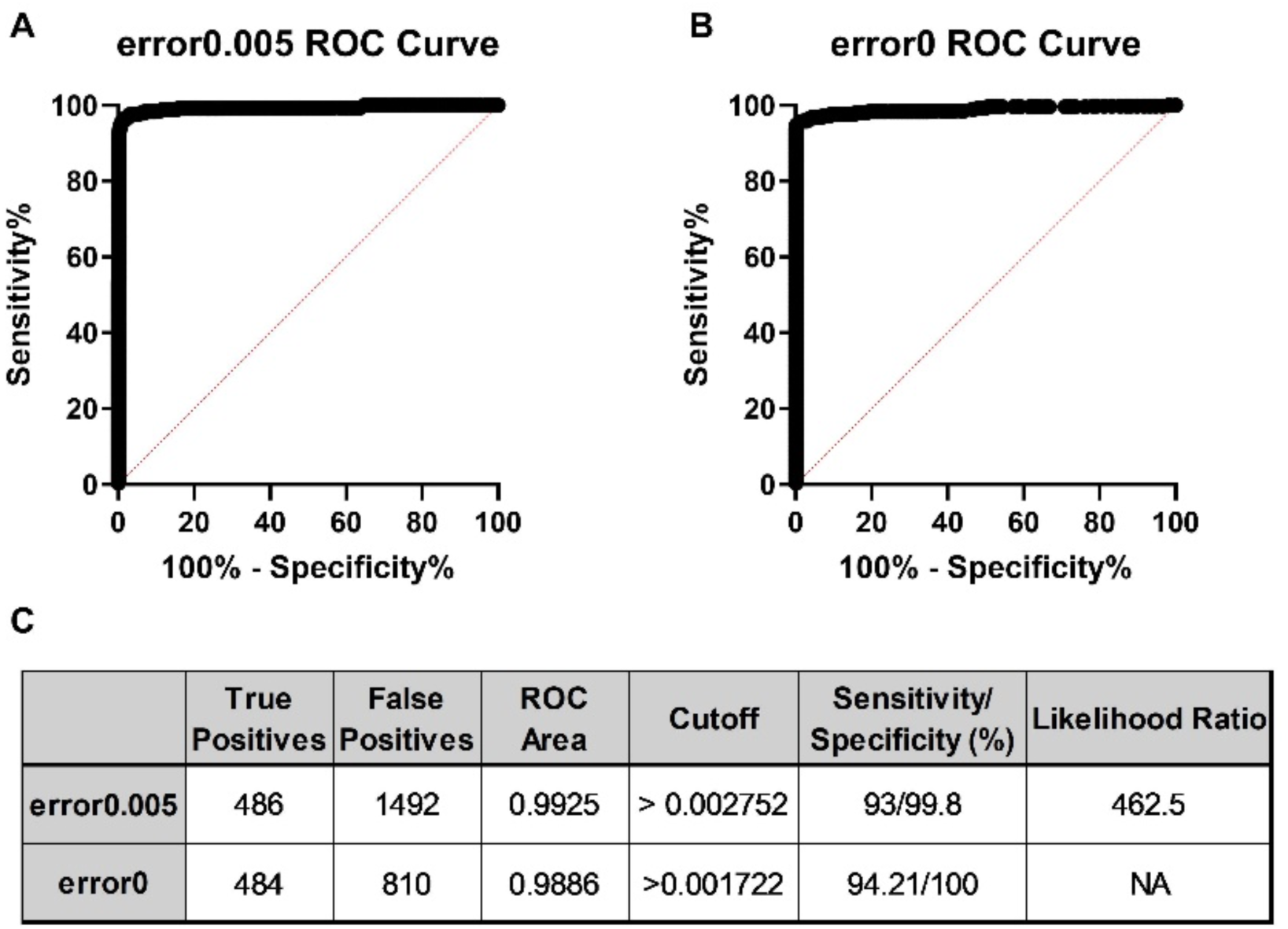
The *wzi*-Seq technique is highly sensitive and specific for detection of strains in mixed samples (A-B) A total of 23 contrived samples containing between 5-58 unique *wzi* types were used as input for the receiver operating characteristic (ROC) analysis using abundance data (readfrac) from both the error0 and error0.005 pipelines. (C) The total number of true positive and false positive values used to generate the ROC curve and ROC curve area is displayed. Using the ROC analysis, readfrac cutoffs with maximum specificity while retaining high sensitivity were chosen for both the error0.005 and error0 pipelines.

### Using wzi-Seq to Investigate Klebsiella Colonization Dynamics

To determine the frequency of mono- or mixed colonization, we analyzed a previously constructed case-control cohort of *Klebsiella-*colonized patients, where cases developed an extraintestinal infection and controls did not^13,20^. A total of 236 rectal swabs were subjected to *wzi*-Seq to determine the frequency of mono- or mixed colonization. Strict criteria for strain detection were applied, including that the abundance must be >1% and that *wzi*-Seq results had to detect *wzi* types previously found by culture and Sanger sequencing of individual isolates from the same rectal swab (see methods). Out of 236 rectal swabs, 27/236 (11.4%) failed *wzi* PCR, 28/236 (11.9%) failed quality control (see methods), 120/236 (50.8%) samples required a second round of *wzi* PCR to meet the DNA input requirements for ONT sequencing, and 10/236 (4.2%) were considered inconclusive. A total of 171 (72.5%) samples passed all quality control metrics and were included in the final analysis. Of these 171 samples, 108/171 (63.2%) were mono-colonized with only one *wzi* type detected and 63/171 (36.8%) were considered mixed with 2-7 *wzi* types present. This contrasts with only 5/171 (2.9%) samples detected as mixed by conventional culture and *wzi* typing of 3 isolates per rectal swab. The majority of mixed samples (79.4%) were colonized with 2 or 3 unique *wzi* types (Fig. 4). Across 171 mono- and mixed colonized samples, 160 unique *wzi* types were identified.

**Figure 4:**
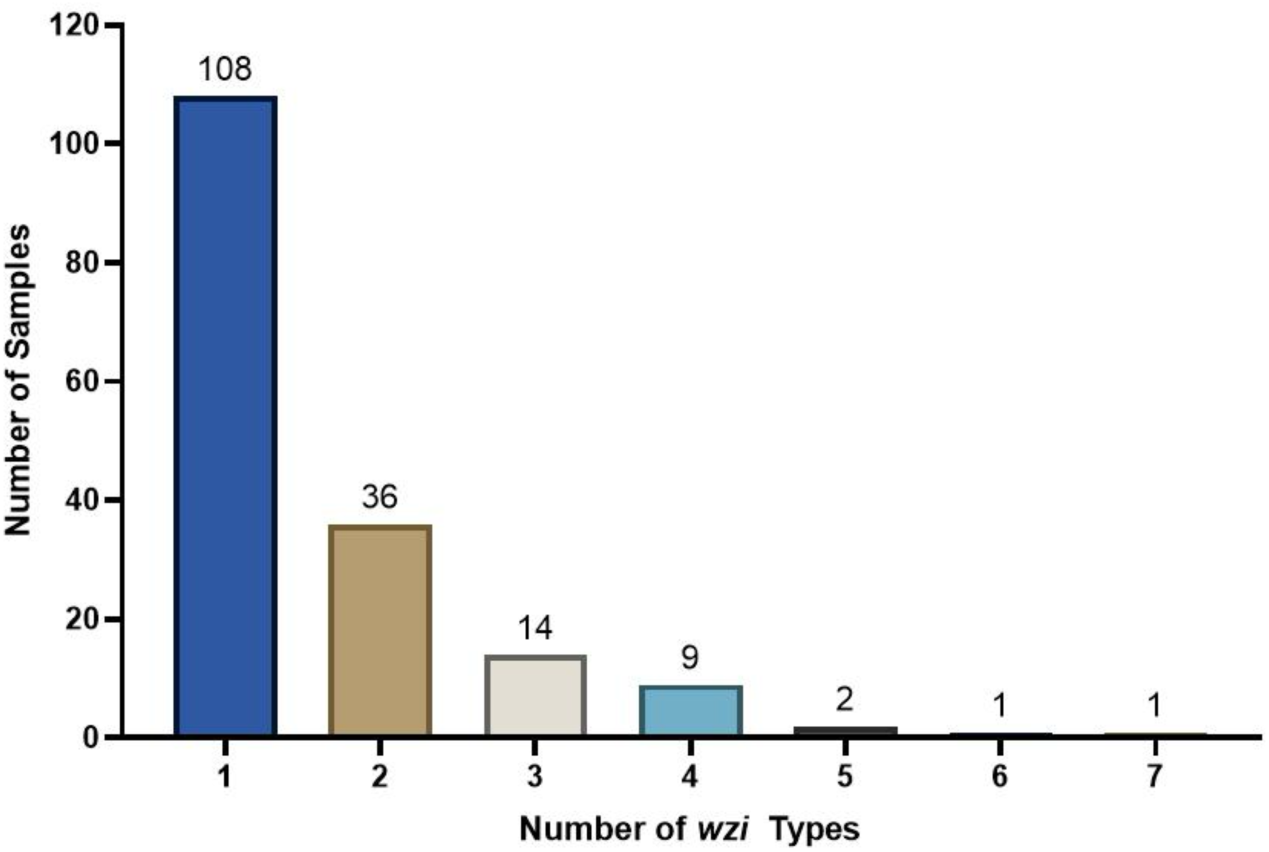
Mono-colonization is more prevalent than mixed colonization in patient rectal swabs The *wzi*-Seq method was used to detect the number of unique *wzi* types present in *Klebsiella* colonized patients from a previously published nested case-control study. Mono-colonization was more prevalent (63.2%) than mixed colonization (36.8%) with mixed samples having been 2-7 unique different *wzi* types.

To further investigate the colonization dynamics in patients colonized with multiple *wzi* types, the abundances of *wzi* types were investigated in mixed samples (Fig. 5). The most abundant of the *wzi* types present (*wzi* Rank 1) had an abundance greater than 50% in 58/63 (92.1%) and greater than 75% in 42/63 (66.7%) of samples (Fig. 6A), indicating that even in mixed samples one *wzi* type tended to dominate (Fig. 6A). Interestingly, there was high diversity in the *wzi* types detected in mixed samples, with 119 unique *wzi* types detected (Fig. 6B,C). There was not a single *wzi* type that tended to be the dominant colonizer, as there were 50 unique *wzi* types detected across mixed samples as the highest *wzi* type (*wzi* Rank 1), and dominant strains in one sample were subdominant in another (e.g. *wzi* 84 in pink, Fig. 6C).

**Figure 5:**
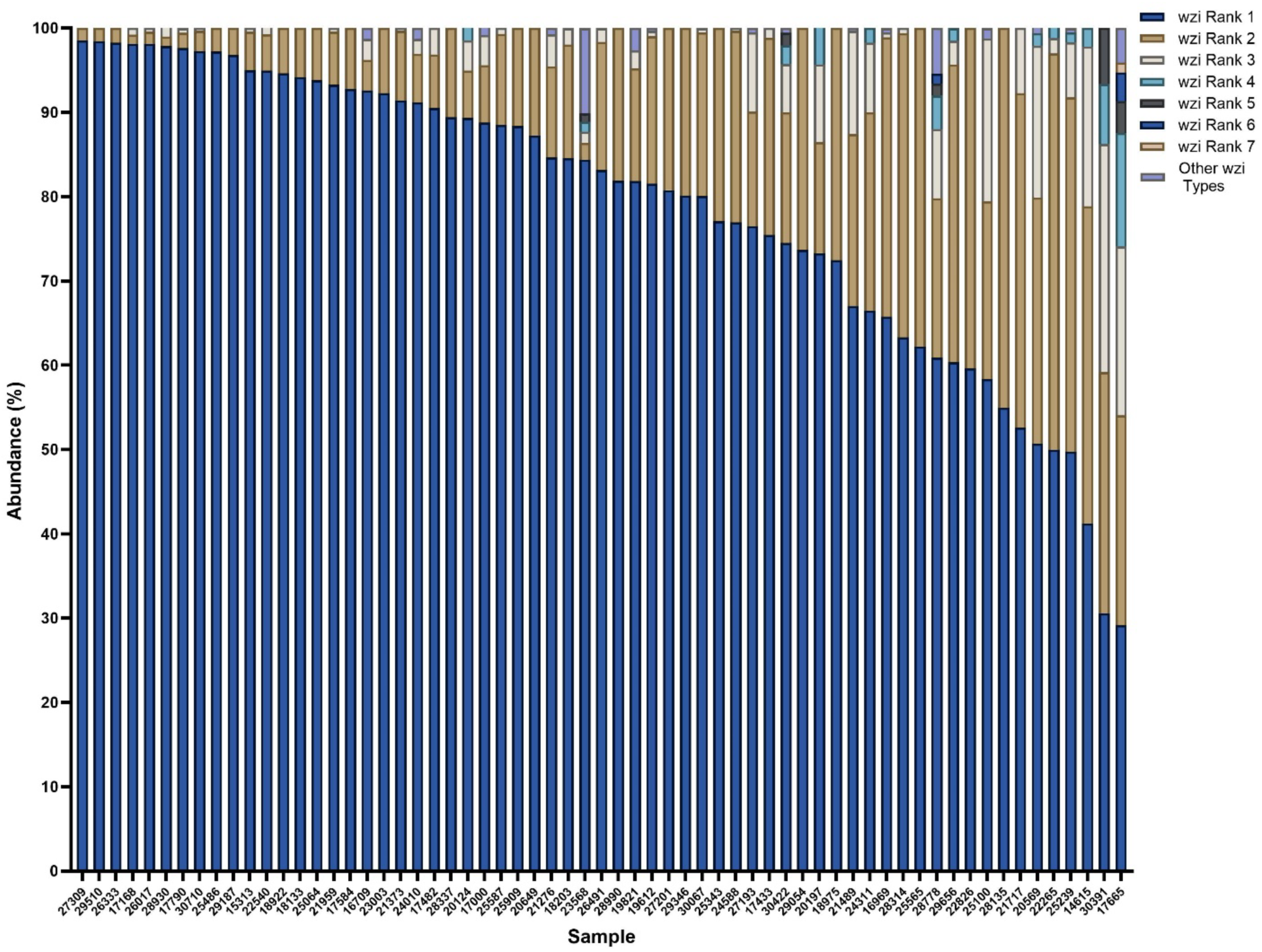
Mixed colonized patient samples are commonly dominated by one *wzi* type The abundances of *wzi* types identified in all mixed samples were quantified with *wzi* rank 1 being the *wzi* type highest in abundance in each sample and *wzi* rank 7 being the lowest in each sample. All *wzi* types with an abundance less than 1% were labeled as “other”.

**Figure 6:**
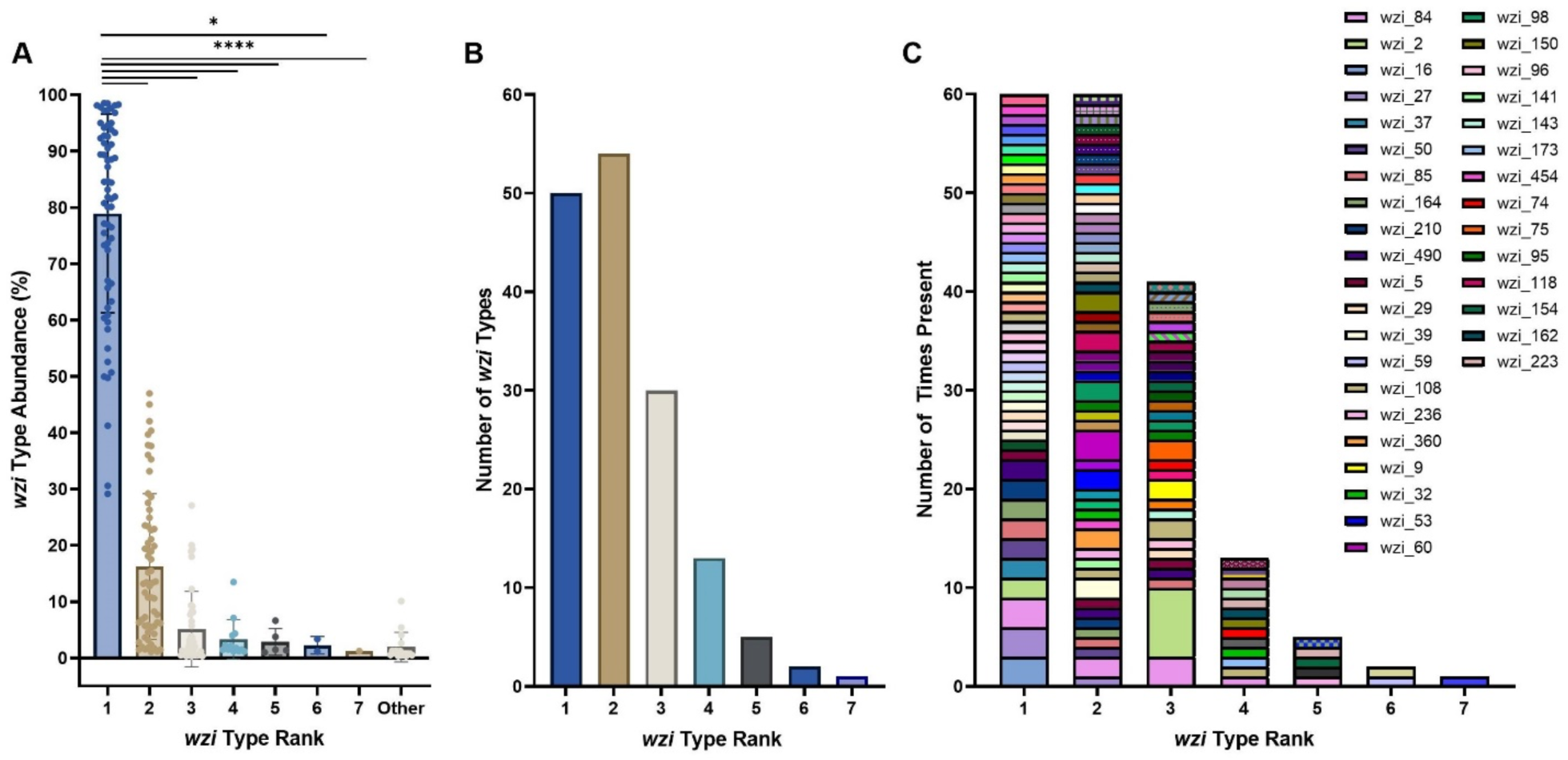
Mixed colonized patients are colonized by diverse *wzi* types The abundance of each *wzi* type based on the abundance ranking was determined. Across all samples, the *wzi* types in rank 1 had significantly higher abundances than any of the other detected *wzi* types, highlighting that mixed samples were uneven with a single *wzi* type dominating samples (A). Across all samples the number of unique *wzi* types present based on the abundance ranking was determined (B). The number of times each unique *wzi* type was present across all samples was quantified in order of the *wzi* type highest to lowest in abundance across all samples (C). The legend displays only colors for *wzi* types that were present more than once in each category. For (A), *, p< 0.05; ****, p<0.0001, by Kruskal-Wallis t-test and n=63.

### Validation of Rectal Swab wzi-Seq Results

To confirm the detection of mixed colonization, repeatability and agreement with an orthogonal method were assessed. The *wzi*-Seq analysis was repeated on a separate sample aliquot and compared to direct Illumina sequencing from the same aliquot. Eight swab samples, four mixed and four mono-colonized based on analysis above, were struck out on MacConkey agar to select for Gram-negative bacteria including *Klebsiella* and then DNA was extracted from plate sweeps for parallel analysis. The repeat *wzi*-Seq analysis reproduced the categorization of mixed and mono-colonization, excluding one sample that failed sequencing (Suppl. Table 5). Data from direct Illumina sequencing of plate sweeps detected *Klebsiella* species in each sample except for 2 samples (Suppl. Fig. 1). For samples with *Klebsiella* detected, reads were mapped to the *wzi-1* reference sequence and *wzi* types were inferred to be present in the population using a popANI approach (see methods). Overall, the results were highly consistent with the rectal swab and plate sweep, with all error0.005 *wzi* types detected, and the same categorization of mixed and mono-colonization (Suppl. Table 5).

Some *Escherichia coli* (*E. coli)* strains can have a *Klebsiella* like capsule and be amplified with *wzi* primers^21,22^. To determine whether *wzi*-Seq was impacted by non-specific amplification of *E. coli wzi* we performed additional bioinformatic analysis. We first evaluated the nucleotide similarity of all 741 unique *Klebsiella wzi* types against all known *E. coli* genomes in the NCBI database using BLAST. Out of all 741 *wzi* types, 6 *wzi* types had 100% identity to known *E. coli* genomes. *E.coli* DNA sequences that had significant but <100% match with any of the 741 *Klebsiella* reference *wzi* types were added to our existing *wzi* reference database. Then the *wzi*-Seq analysis pipeline using the updated reference database was rerun on rectal swabs with mixed colonization (see methods). Out of 63 mixed samples, 62/63 (98.4%) were still classified as mixed after running the *E. coli* amended pipeline; one had a second *Klebsiella*-specific *wzi* type that fell to 0.9%, just below our 1% cutoff for a mixed sample. Five samples contained at least one *wzi* type (wzi_5, wzi_370, or wzi_173) that was 100% identical to an *E. coli wzi* type, but this did not change the result of mixed detection because either this *wzi* type was from a culture-confirmed *K. pneumoniae* isolate or there were multiple *Klebsiella*-specific *wzi* types present in the sample (Supplemental Dataset).

### Mono-colonization is Associated with an Increased Risk of Klebsiella Infection

To determine whether mono- or mixed colonization was associated with an increased risk of developing a *Klebsiella* infection, patient characteristics associated with each rectal swab were evaluated. Out of 171 rectal swabs, 66 were from colonized patients who developed a subsequent *Klebsiella* infection (case) while 105 were from asymptomatic colonized controls (Suppl. Fig. 2A). Mixed colonization was more prevalent in control samples (45/105, 43%) than in cases (18/66, 27%) (Suppl. Fig. 2A, B).

*Klebsiella* dominance in the gut is associated with an increased risk of infection, so we compared *Klebsiella* abundance in mono- and mixed colonization as previously measured by our validated qPCR assay^7,8,11^. *Klebsiella* relative abundance in mono-colonized samples was significantly higher in cases than controls but this was not the case in mixed colonized samples (Suppl. Fig. 3A, B). Overall, the relative abundance was significantly higher in mono-than in mixed colonized patients (Suppl. Fig. 3C).

To determine the association of mono- and mixed colonization with subsequent infection, we constructed a multivariable model. Patient characteristics could be putative confounders to consider for adjustment (see methods), so we characterized the distribution of these variables versus subsequent infection. Table 2 shows important baseline characteristics of the study population and variables that were considered in our final model. Relative abundance and mono/mixed colonization status were found to be colinear. If they lie along the same causal pathway and with the observed collinearity, then including *both* relative abundance and mono-colonization in the model together would result in weakening the association. Indeed, we observed that while both mono-colonization and relative abundance are both associated with infection (Table 2), including both as covariates weakens the association with infection and there is no interaction noted statistically (p=0.691 for the interaction vs. infection in a multiple logit model). Therefore, relative abundance was not considered for the final model, and we focused on mono-colonization vs. mixed colonization^11,13^. Other than age, gender and collection month, which were *a priori* factors to include for adjustment (see methods), only prior urinary catheter use and baseline serum albumin were retained in the final model. After adjustment for these potential confounders (Suppl. Table 6) we measured an independent association between mono-colonization and subsequent infection demonstrating a 2.18-fold increased odds of infection (95% CI 1.08-4.57; p=0.034).

**Table 2:** Case-control study population characteristics

| Characteristic | No infection<sup>1</sup> | Infection<sup>1</sup> | p-value<sup>2</sup> |
| --- | --- | --- | --- |
|  | N = 105 | N = 66 |  |
| **Age<sup>3</sup>** | 61 (54, 70) | 59 (52, 69) | 0.6 |
| **Gender<sup>3</sup>** |  |  | >0.9 |
| Female | 49 (47%) | 31 (47%) |  |
| Male | 56 (53%) | 35 (53%) |  |
| **Race** |  |  | 0.8 |
| Non-white | 21 (20%) | 12 (18%) |  |
| White | 84 (80%) | 54 (82%) |  |
| **Colonization<sup>3</sup>** |  |  | 0.04 |
| Mixed | 45 (43%) | 18 (27%) |  |
| Mono | 60 (57%) | 48 (73%) |  |
| ***Klebsiella* relative abundance** | 4 (0, 20) | 20 (1, 59) | 0.002 |
| **Weighted Elixhauser Score** | 19 (12, 26) | 23 (16, 28) | 0.06 |
| **Depression** | 25 (24%) | 25 (38%) | 0.049 |
| **Prior/Baseline Urinary Catheter<sup>3</sup>** |  |  | 0.006 |
| FALSE | 49 (47%) | 17 (26%) |  |
| TRUE | 56 (53%) | 49 (74%) |  |
| **Prior/Baseline Feeding Tube** |  |  | 0.056 |
| FALSE | 103 (98%) | 60 (91%) |  |
| TRUE | 2 (1.9%) | 6 (9.1%) |  |
| **Prior Broad Spectrum Antibiotic Exposure** | 19 (18%) | 22 (33%) | 0.023 |
| **Prior/Baseline Pressor Dependent Hypotension** | 11 (10%) | 15 (23%) | 0.03 |
| **Baseline Serum Albumin<sup>3</sup>** |  |  | 0.003 |
| <2.5 | 21 (20%) | 29 (44%) |  |
| ≥2.5 | 79 (75%) | 34 (52%) |  |
| NA | 5 (4.8%) | 3 (4.5%) |  |
| <sup>1</sup>Results reported as median (Q1, Q3) or n (%) as appropriate |  |  |  |
| <sup>2</sup>Wilcoxon rank sum test, Pearson's Chi-squared test, or Fisher's exact test as appropriate |  |  |  |
| <sup>3</sup>These variables were included in the final multivariable model. |  |  |  |

### Gut Community Structures Do Not Vary Between Mono or Mixed Colonized Patients

To investigate whether overall microbial community differences contribute to *Klebsiella* colonization dynamics, we analyzed available 16S rRNA gene sequencing of the patient rectal swabs^23^. Interestingly, the gut microbial community structure was not significantly different between mono- or mixed colonized patients. No significant differences in community richness, evenness, or diversity were observed between patient groups (Fig. 7A-C). Furthermore, a principal coordinate analysis did not reveal any significant differences in the beta-diversity between mono- and mixed colonized patients (Fig. 7D). When using 16S rRNA data to assess *Klebsiella* abundance, higher abundance of the ASV: ASV000001 *Klebsiella* was observed in mono compared to mixed samples that approached statistical significance (p=0.06; Fig. 7E), consistent with our qPCR data.

**Figure 7:**
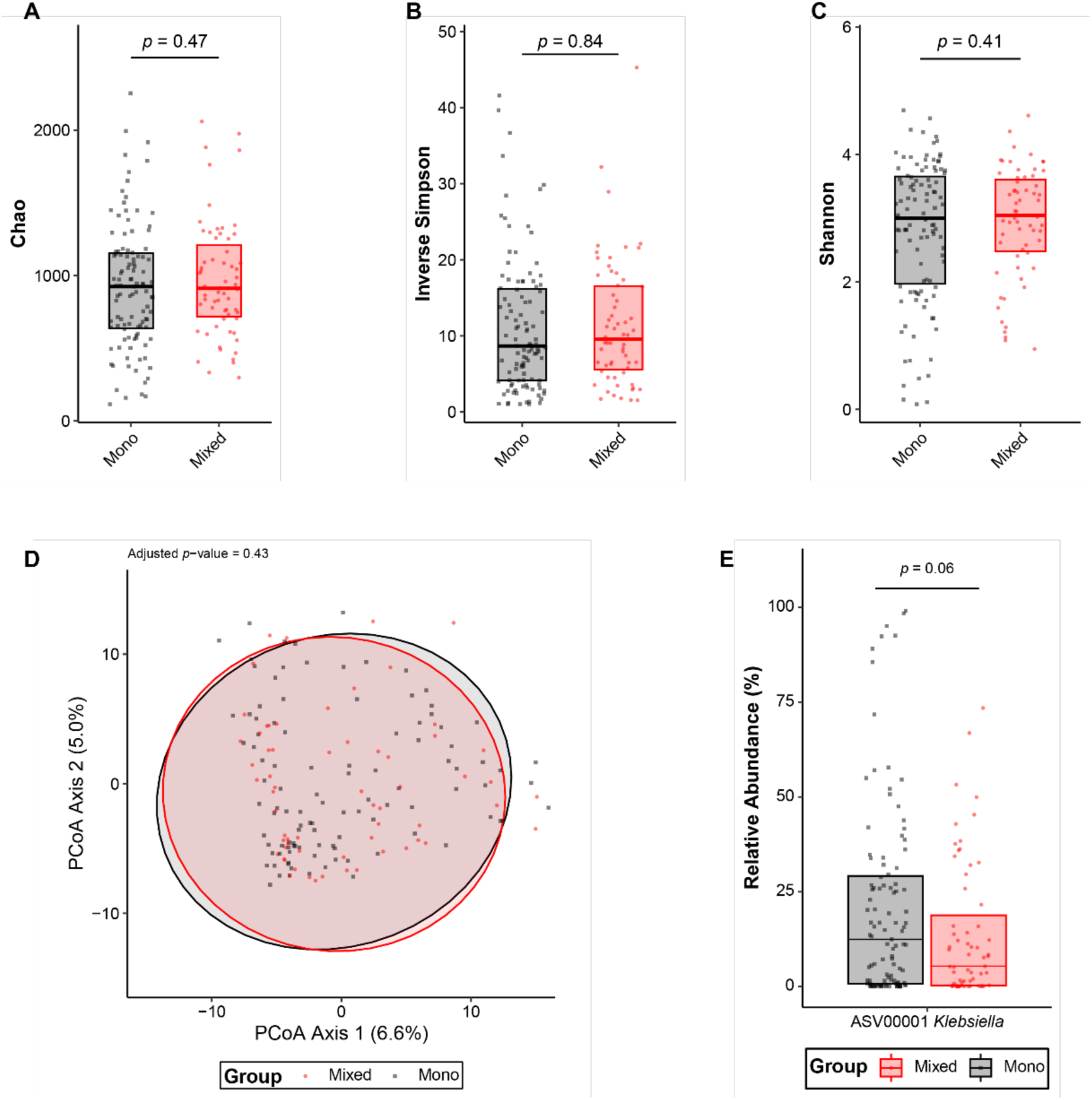
Gut community structures do not differ significantly between mono- or mixed colonized patients The gut community structure using alpha (A-C) and beta (D) diversity metrics were assessed via 16S sequencing of patient rectal swabs and the bioinformatic package, mothur. No significant differences were in alpha diversity metrics including Chao (A), inverse Simpson (B), or Shannon were observed (C). There were no significant differences in beta diversity (D) between mono- or mixed colonized patients. Based on ASVs higher relative abundance levels (E) were observed in mono-compared to mixed colonized patients but this did not reach statistical significance. For all analyses, a p-value ≤ 0.05, after Benjamini-Hochberg adjustment where appropriate was considered significant.

## Discussion

In this study, we describe the development and validation of *wzi*-Seq, a novel amplicon-based sequencing method to measure *Kp* population structure in samples (Fig. 1). We demonstrate that the *wzi*-Seq method has excellent read accuracy and high precision for measuring relative abundance, and these performance characteristics are not impacted when performing additional rounds of PCR (Fig. 2; Table 1; Suppl. Tables 1-3). Furthermore, the assay has a sensitivity and specificity of 93/99.8% for the error0.005 pipeline and 94.21/100% for the error0 pipeline for strain detection in samples with up to 58 strains (Fig. 3; Suppl. Table 4). Applying *wzi*-Seq to a cohort of *Kp* colonized patients, we determined that *Kp* mono-colonization was more prevalent than mixed colonization, with mono-colonization being independently associated with an increased infection risk (OR=2.18; p=0.034). This study describes a novel and reliable technique to investigate *Kp* colonization dynamics and reveals the population structure of colonizing strains as a potentially important risk factor for infection.

The *wzi*-Seq assay has performance characteristics that make it suitable for several applications. This amplicon sequencing approach can perform strain typing directly from samples, distinguishing it from WGS, MLST, PGEF, and MALDI-TOF that typically require culturing of single *Klebsiella* isolates. Importantly, *wzi* typing has similar discriminatory power as MLST and K-typing^7,16,24^. Also, the assay can accurately detect and quantify mixtures of *Kp* strains within a sample, similarly to 16S rRNA gene sequencing for multiple species. We developed the *wzi*-Seq assay using nanopore sequencing in part because the long region of interest precluded Illumina sequencing. We have taken advantage of recent advances in nanopore accuracy at both the sequencing and base calling steps, and mitigated the residual error rate by using read mapping to known sequences and applying several quality filtering steps^25^. As used here, this method can be used to detect and quantify multiple strains in a sample with high confidence.

The finding that mixed colonization occurs in a substantial subset of patients, but that mono-colonization is associated with increased infection risk has several implications. Mixed colonization and infection has been documented in other pathogens including *Clostridioides difficile*^26–29^. Unlike *Klebsiella,* C. *difficile* causes intestinal disease and mixed infection with multiple strains was significantly associated (OR 3.5, 95% CI 1.3-9.4, p = 0.015) with recurrence in patients, even after adjustment for gender and previous exposure to antibiotics^26^. For *Klebsiella*, mixed colonization may represent a more commensal state than mono-colonization. Indeed, a recent study illustrated that two strains of *Kp* can cross-feed one another with one strain compensating for the fact that the other cannot utilize a specific nutrient^30,31^. This may represent a stable colonization state where one strain is kept in check by its reliance on the other, reducing the chance that they cause clinical disease. In contrast, mono-colonization was associated with higher relative abundance of *Kp*, and these variables were co-linear in our models of infection. This may represent the outcome of competition with other *Kp* strains, resulting in complete displacement of one strain and replacement with the other. Even in mixed colonization, one *wzi* type tended to dominate, suggesting that *Kp* strains outcompete other strains, or at least settle at a higher density in equilibrium. It would be interesting to explore whether mixed colonized patients are at higher risk for prolonged colonization, or whether over time these patients progress to a mono-colonized state due to dominance by a single strain.

Furthermore, this highlights that typing only single isolates may miss these mixed colonized states with implications not only on patient infection risk but also potential transmission events.

We also found that the gut microbial community was not significantly different between mono-and mixed colonized patients, suggesting that microbial richness or diversity in the gut does not impact strain colonization status (Fig. 7). This is consistent with previous studies that only observed minor changes in the gut microbial community during *Kp* colonization in mice^32–34^. This suggests that in our patient cohort, other clinical and environmental factors might play a more significant role in shaping the overall community structure. Alternatively, *Kp* strains might occupy a particular ecological niche within the gut microbiome that allows coexistence with other microbes without causing substantial shifts in the community structure.

This study and the *wzi*-Seq method have several limitations. As *wzi*-Seq relies on a single conserved gene for discrimination, the technique cannot be used to differentiate between strains of the same *wzi* type. Furthermore, the primers used to amplify the *wzi* gene can, in rare cases, amplify group 1 *E. coli wzi* genes^21,22^. The incidence of *Klebsiella* capsules in *E. coli* is unclear, although a study of freshwater *E. coli* found *Klebsiella* capsules in only 7% of isolates^35^. We addressed this limitation by including *E. coli wzi* sequences in our reference database for read-mapping and requiring a 100% match to a *Klebsiella wzi* sequence to consider that *wzi* type to be present in a given sample. When using complex samples with unknown *wzi* types, orthogonal validation approaches may be needed to distinguish *Kp* and *E. coli* with identical *wzi* sequences. In conclusion, this study describes the development and validation of *wzi*-Seq, a novel technique that enables the detection and quantification of *Kp* from complex samples. In future directions, the *wzi*-Seq technique can be applied to test patient samples, adapted to target additional genes, and applied to characterize diverse populations of *Kp* in both experimental and natural conditions.

## Methods

### Rectal Swab Sample Collection and Analysis

Patient enrollment and sample collection was performed previously and was approved by the Institutional Review Board^13,20^. Patient rectal swabs were collected with instructions to pass the anal verge using flocked swabs placed in Amies transport media (Becton, Dickinson, Franklin Lakes, NJ), and screened for *Klebsiella* via plating on MacConkey agar and verification via MALDI TOF mass spectrometry^13^. From positive rectal swabs, up to three *Klebsiella* isolates were collected and subject to *wzi* PCR and Sanger sequencing and the *wzi* type of each isolate was determined by matching the consensus *wzi* sequence to the BigsDB database (https://bigsdb.pasteur.fr/klebsiella/klebsiella.html).

### *wzi*-Seq Assay

DNA was extracted from rectal swabs or contrived samples using the Qiagen UltraClean Microbial DNA kit (Qiagen, Hilden, Germany) as per the manufacturer’s instructions. PCR was performed using previously published *wzi* primers and the Q5 polymerase (New England Biolabs, Ipswich, MA, USA) with volumes as specified by the manufacturer in a 100 µL reaction^16^. For all reactions, 4 µL of DNA template with a concentration of 1-2ng/µL was used. PCR conditions were 30 sec at 98°C, then 30 cycles of 10 sec at 98°C, 30 sec at 59°C, 30 sec at 72°C, followed by 2 minutes at 72°C on a MiniAmp Plus Thermal Cycler (Thermo Fisher Scientific, Waltham, MA, USA). For samples with poor amplification a second round of PCR was performed using the same conditions as listed above, using 4µL of the first PCR reaction as the template. Samples were run on a 1% agarose gel at 100V for 30 minutes to confirm amplification, purified using the QIAquick PCR Purification Kit (Qiagen, Hilden, Germany), and quantified using Nanodrop and Qubit (Thermo Fisher Scientific, Waltham, MA, USA). All samples were submitted to the University of Michigan Advanced Genomics Core (AGC) for Oxford Nanopore Technologies sequencing (Oxford Nanopore Technology, Oxford, UK). Basecalling was performed using the SUP pipeline by the University of Michigan AGC.

### Bioinformatic analysis

#### error0 and error0.005 Pipelines

Reads with mean base quality lower than 10, or with more than 5% bases lower than 10 were removed using NanoFilt v2.8.0^36^ and a custom script. Remaining reads were aligned against the reference *wzi* types curated from the BIGSdb-Pasteur database (https://bigsdb.pasteur.fr/klebsiella/) using Minimap2 v2.26^37^. Reads were filtered by these criteria sequentially: mapping quality was larger than zero; the read had a single mapping location after mapping quality filtering; alignment covered more than 95% of the mapped gene sequence; NM value (number of mismatches and gaps) was zero. Filtered reads were counted on each reference gene sequence.

The filtering for the error0.005 pipeline was the same as the error0 pipeline except that the base quality fraction filter was disabled, and the error rate filter was relaxed to lower than 0.5% of the alignment length.

#### Quality criteria for rectal swab samples

To ensure confidence in identified *wzi* types the following quality control metrics were applied to all samples: 1) error0.005 samples had to have greater than 1,000 reads, 2) error0.005 and error0 readfrac cutoffs (>0.00275 and >0.00175, respectively) were applied before abundance calculations, 3) samples had to detect *wzi* types from banked rectal isolates from that sample (168/171 available), and 4) *wzi* type abundance had to be greater than 1%. A 1% abundance cutoff was applied as this was the lowest abundance validated using contrived mixtures. To reduce the possibility of contamination causing the appearance of mixed colonization, samples were considered inconclusive and not included if the wzi_2 allele of our commonly used laboratory strain was present as the first or second most abundant *wzi* type and was not detected via rectal isolate typing. Samples were then examined for concordance between error0.005 and error0 pipelines. If there was discordance, BAM files were examined in IGV for potential single nucleotide polymorphisms that may explain the discordance and where possible, checked against sequenced genomes from that specimen.

#### Detection of potential E. coli capsule sequences

To map reads to a reference database with both *Klebsiella* and potential *E. coli wzi* alleles, the error0 pipeline was run with the following changes: 1) the mapping quality filter was removed, 2) zero mismatches were allowed between mapped reads and the reference, 3) 100% coverage of the reference was required, and 4) a unique mapping position needed to remain after steps 2 and 3. Mixed samples were re-run using this *E. coli* amended pipeline to determine whether this impacted the mixed status of any samples.

### Illumina Sequencing Analysis

#### Data QC and determination of species composition

Fastqc (v0.12.1) (https://www.bioinformatics.babraham.ac.uk/projects/fastqc/) was used to assess the quality of sequencing reads before and after trimming with Trimmomatic (v0.36)^38^. InStrain (v1.10.0) was used to identify bacterial species composition for each population sample^39^. The reference database consisted of 4,644 representative genomes from the Unified Human Gastrointestinal Genome (UHGG) collection in 2025. Trimmed reads were aligned to each reference independently with Bowtie2 (v2.4.2). inStrain output was filtered with 50% genome breadth cutoff.

#### wzi popANI pipeline

BWA mem (v0.7.17) (https://arxiv.org/abs/1303.3997), was used to align trimmed reads to *wzi-1*. Samtools (v 1.21) was used to transform sam files into bam files and index the bam files^40^. R (v4.4.3) was used to calculate the popANI between population sequences and wzi-references, which provides the lower-bound distance to each *wzi* type. The value of popANI to a given wzi-type increases if there are no shared alleles between a sample and *wzi* reference at a given position in the alignment. For each sample, *wzi* types were removed if popANI was >3 or higher than the lowest popANI value. Breadth of coverage was estimated for each remaining *wzi* type and *wzi* types were removed if breadth of coverage was lower than 99%.

### 16S rRNA sequencing and analysis

DNA extraction and 16S rRNA gene sequencing was performed elsewhere^23^. Sequences were processed with mothur (v. 1.48.0) and aligned to the SILVA reference alignment, release 132^41,42^. The sequencing error rate was assessed using a predefined mock community and estimated to be 0.0066%. For all analyses, sample read counts were rarefied to 8,363 reads (lowest sample read count >8,000), and one sample (PR24765, prokka_sample_923-JV-132, rectal swab corresponding to patient colonized by *Klebsiella* strain Kp7994) was excluded due to low total read count (<8,000). Differences in community structure were assessed by PERMANOVA (1,000 permutations) using robust Aitchison distance as the dissimilarity metric from the vegan package, v. 2.6-2, and alpha diversity metrics were calculated using mothur^41,43^. Data analysis was carried out in RStudio 2021.09.0+351 “Ghost Orchid” Release for macOS. For all analyses, a p-value ≤ 0.05, after Benjamini-Hochberg adjustment where appropriate, was considered statistically significant. All scripts for 16S rRNA gene sequencing analysis used for this study are available at https://github.com/jayvorn/wzi_mixed_mono_colonization (embargoed until publication).

### Logistic Regression Model and Analysis

The goal of this analysis was to determine the independent contribution of mono-colonization vs. mixed colonization with *Kp* to risk of subsequent clinical infection from *Kp*. We did this by including mono-colonization as the primary predictor and then included covariates due to a priori importance (e.g., past matching criteria) or based on retention as putative confounders after a stepwise selection procedure. For the former types of covariates, our prior studies on the same cohort from which the current study’s subjects were drawn, informed the clinical variables to consider for adjustment^20^. Notably, since our source cohort was from a matched case control study but not all cases and controls could be included here, we included the matching criteria of age, gender, and month of swab collection as covariates for adjustment in our model^13^. We then turned to purposeful selection to determine which other putative confounders to include in the final model^44^. Starting with variables capturing demographics, comorbidities, prior/baseline medication and device exposures, and prior laboratory parameters, we proceeded in a stepwise fashion until only significant variables and potential confounders were retained, and this is the final model presented in the results^44^.

## Data availability

The 16S sequencing data used in this study are deposited in the Sequence Read Archive (SRA) database under accession code PRJNA789565. The *wzi*-sequencing data will be deposited and available in SRA prior to publication. Patient variables used in logistic regression can be made available upon request, after ensuring de-identification and compliance with IRB regulations at both institutions.

## Funding Statement

Research reported in this publication was supported by the National Institute of Allergy and Infectious Diseases of the National Institutes of Health under Award Number R21AI176642. The content is solely the responsibility of the authors and does not necessarily represent the official views of the National Institutes of Health.

## Competing Interests

Dr. Vornhagen has consulted for Vedanta Biosciences, Inc. Dr. Rao has received an investigator-initiated grant from Merck & Co, Inc.; he has consulted for Seres Therapeutics, Inc., Rebiotix, Inc., Vedanta Biosciences, Inc., and Summit Therapeutics, Inc.

---

**Supplemental Figure 1:**
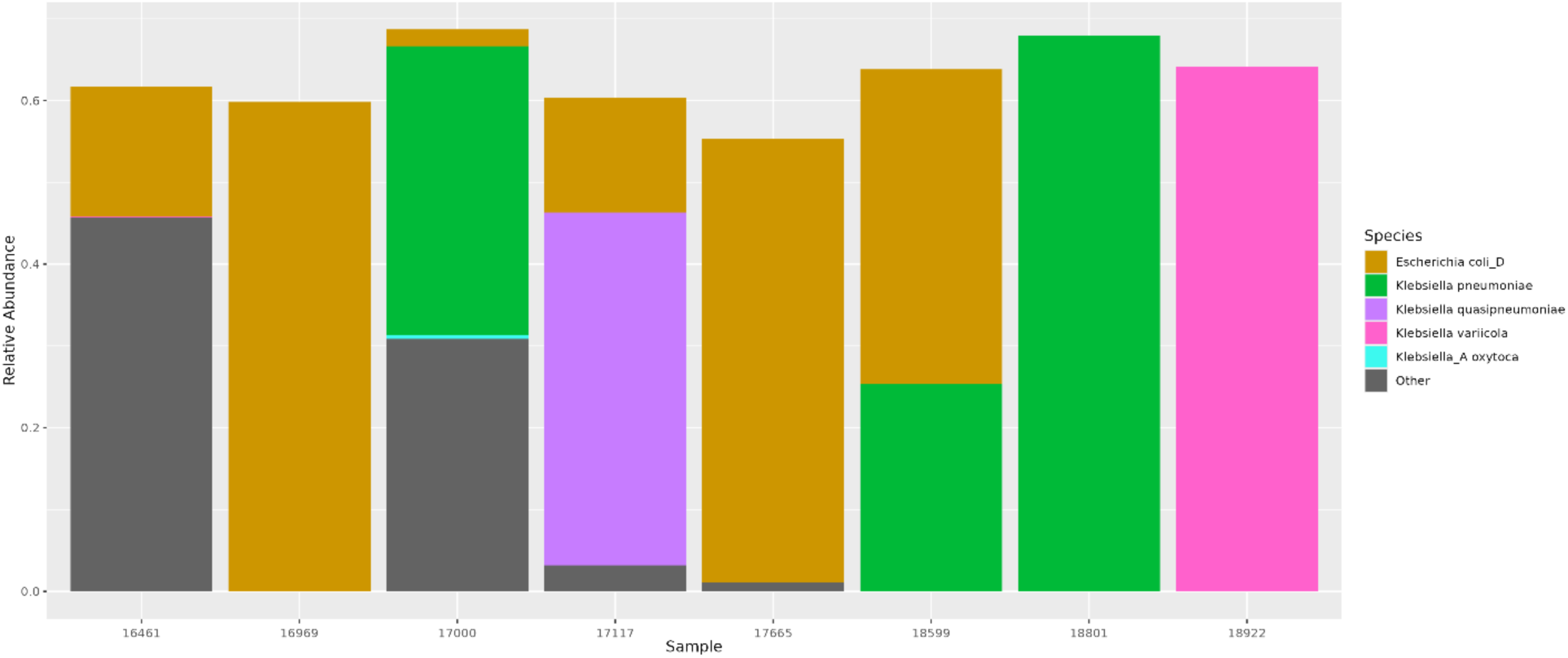
Species composition based on Illumina Sequencing of *Klebsiella* Mixed and Mono-Colonized Rectal Swabs used for validation. Eight glycerol amended swab samples, four mixed and four mono-colonized based on analysis above, were struck out on MacConkey agar to select for Gram-negative and then DNA was extracted. Samples were subjected to Illumina sequencing and bacterial species composition was determined by InStrain.

**Supplemental Figure 2:**
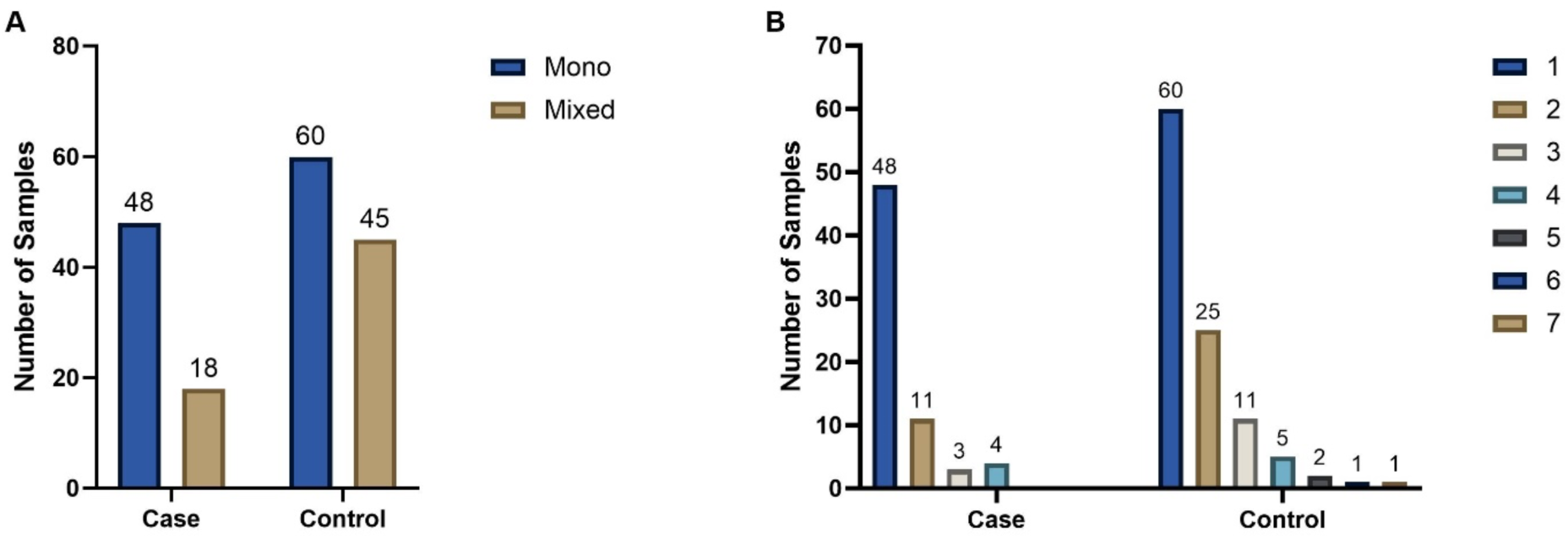
Mono- and mixed colonization in cases of *Klebsiella* infection and controls Colonization and case status of all patients were evaluated based on mono- or mixed colonization status (A) or the number of *wzi* types present in each sample (B).

**Supplemental Figure 3:**
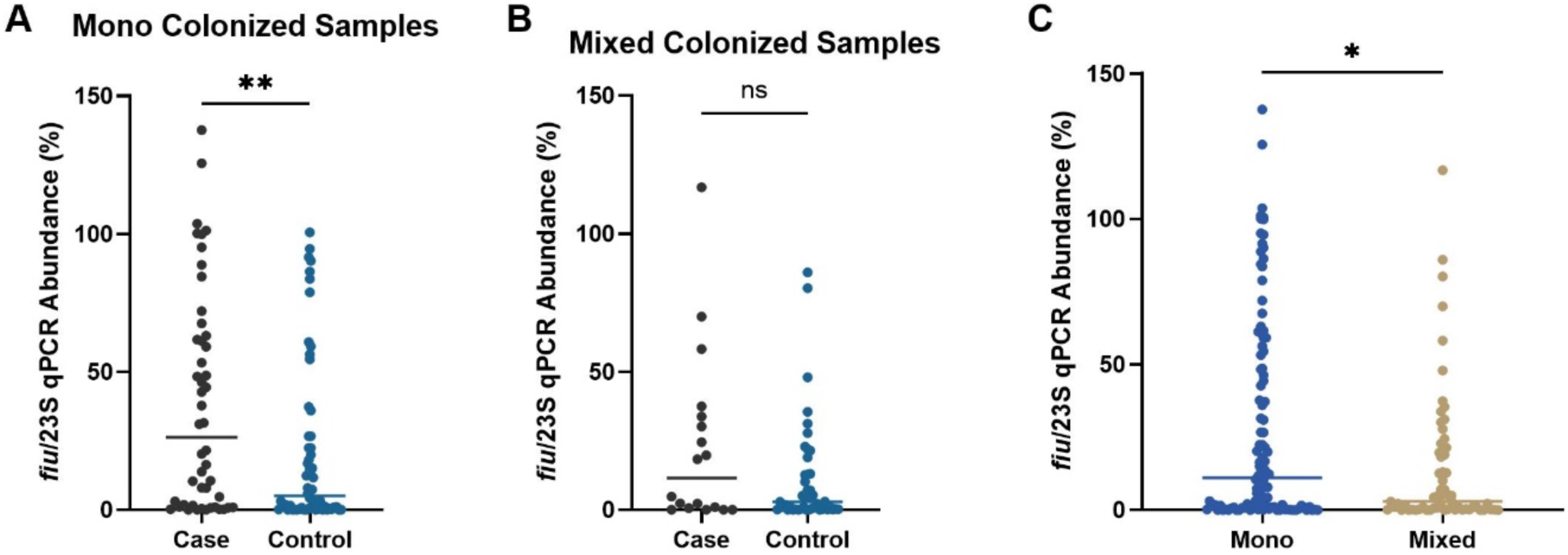
Mono-colonized case samples have significantly higher *Klebsiella* relative abundance than controls (A-C) The *Klebsiella* relative abundance in each rectal swab was quantified using a validated qPCR assay quantifying the conserved *Klebsiella fiu* gene and the panbacterial 23S rRNA gene. The ratio of the *fiu* and 23S rRNA gene were used to calculate the *Klebsiella* relative abundance in case or control patient rectal swabs from mono- (A) or mixed (B) colonized patients. The *Klebsiella* relative abundance calculated previously (11) was also compared between mono-and mixed colonized patients (C). For (A-B), **, p< 0.001 using a Mann-Whitney t-test with n≥18. For (C), *, p< 0.05 with an n=171 and Mann-Whitney t-test.

**Supplemental Table 1:** Applying *wzi-*Seq to a single *Klebsiella* strain sample

| Pipeline | Reads After Filtering | Total Correct Reads | Total Incorrect Reads | Accurate Reads (%) | Total *wzi* types |
| --- | --- | --- | --- | --- | --- |
| error0.005 | 13,367 | 13,338 | 29 | 99.78 | 8 |
| error0 | 6,181 | 6,179 | 2 | 99.96 | 3 |

**Supplemental Table 2:**
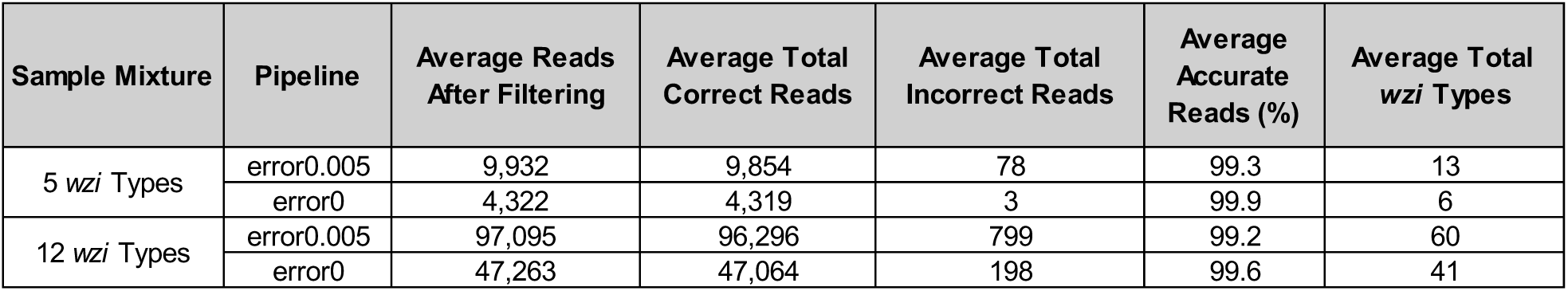
The *wzi*-Seq method has high read accuracy with 5 and 12 *wzi* type mixtures

| Sample Mixture | Pipeline | Average Reads After Filtering | Average Total Correct Reads | Average Total Incorrect Reads | Average Accurate Reads (%) | Average Total *wzi* Types |
| --- | --- | --- | --- | --- | --- | --- |
| 5 *wzi* Types | error0.005 | 9,932 | 9,854 | 78 | 99.3 | 13 |
|  | error0 | 4,322 | 4,319 | 3 | 99.9 | 6 |
| 12 *wzi* Types | error0.005 | 97,095 | 96,296 | 799 | 99.2 | 60 |
|  | error0 | 47,263 | 47,064 | 198 | 99.6 | 41 |

**Supplemental Table 3:**
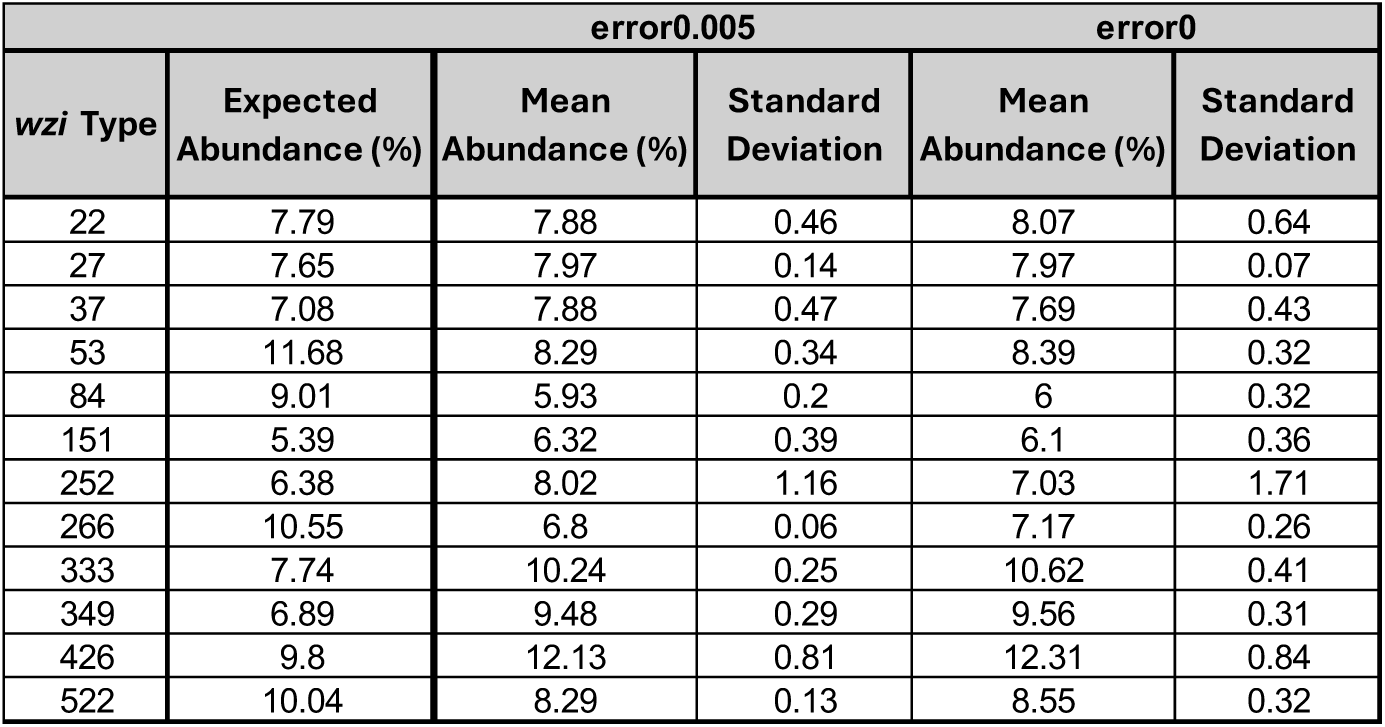
Summary table showing high precision between replicates for the 12 *wzi* type mixture

**Supplemental Table 4:** Contrived 30 and 58 *wzi* type mixtures have decreased accuracy with the error0.005 pipeline

| Sample Mixture | Pipeline | Reads After Filtering | Total Correct Reads | Total Incorrect Reads | Accurate Reads (%) | Total *wzi* Types | Correct *wzi* Types |
| --- | --- | --- | --- | --- | --- | --- | --- |
| 30 *wzi* Types | error0.005 | 58200 | 55562 | 2638 | 95.47 | 99 | 29 |
|  | error0 | 27446 | 27296 | 150 | 99.46 | 56 | 28 |
| 58 *wzi* Types | error0.005 | 39822 | 38637 | 1185 | 97.02 | 114 | 53 |
|  | error0 | 18836 | 18744 | 92 | 99.51 | 76 | 53 |

**Supplemental Table 5:** Illumina Sequencing of Rectal Swab Plate Sweeps Validates *wzi*-Seq Results

| Sample | Rectal Swab Status | Rectal Swab *wzi* Types (>1%) | Plate Sweep Status (*wzi*-Seq) | Plate Sweep *wzi*-Seq *wzi* Types (> 1%) | Plate Sweep Status (Illumina) | Plate Sweep Illumina *wzi* Types |
| --- | --- | --- | --- | --- | --- | --- |
| 16461 | Mono | *wzi* 427 | Mono | *wzi* 427 | Mono | *wzi* 427 |
| 16969 | Mixed | *wzi* 527, *wzi* 27 | Failed | Failed | Failed | Failed |
| 17000 | Mixed | *wzi* 273, *wzi* 300, *wzi* 84 | Mixed | *wzi* 273, *wzi* 300, *wzi* 84 | Mixed | *wzi* 273, *wzi* 300, *wzi* 84 |
| 17117 | Mono | *wzi* 537 | Mono | *wzi* 537 | Mono | *wzi* 537 |
| 17665 | Mixed | *wzi* 360, *wzi* 537, *wzi* 9, *wzi* 108, *wzi* 223, *wzi* 227, *wzi* 340 | Mixed | *wzi* 360, *wzi* 537, *wzi* 9, *wzi* 108, *wzi* 223, *wzi* 227, *wzi* 340, *wzi* 158 |  |  |
| 18599 | Mono | *wzi* 19 | Mono | *wzi* 19 | Mono | *wzi* 19 |
| 18801 | Mono | *wzi* 173 | Mono | *wzi* 173 | Mono | *wzi* 173 |
| 18922 | Mixed | *wzi* 379, *wzi* 53 | Mixed | *wzi* 379, *wzi* 53 | Mixed | *wzi* 379, *wzi* 53 |

**Supplemental Table 6:** Results from final multivariable model of gut mono-colonization vs. mixed colonization and the risk of subsequent *Klebsiella* infection

| Variable | Odds Ratio | 95% CI (Lower - Upper) | P Value |
| --- | --- | --- | --- |
| Age | 0.989 | 0.961 – 1.02 | 0.417 |
| Male Gender | 0.835 | 0.421 – 1.64 | 0.602 |
| Gut Colonization Month<sup>1</sup> | 1.02 | 0.955 – 1.08 | 0.608 |
| Mono-colonization | 2.18 | 1.08 – 4.57 | 0.034 |
| Prior Urinary Catheter | 1.92 | 0.947 – 4.00 | 0.0733 |
| Serum albumin ≥2.5 g/dL | 0.353 | 0.167 – 0.731 | 0.00546 |
| Serum albumin missing | 0.382 | 0.0688 – 1.82 | 0.234 |
| <sup>1</sup>Colonization month was coded from 0-18 chronologically for each month that gut colonized subjects were recruited. |  |  |  |

## Notes

### Summary of Updates

Added supplemental data. Added funding information and competing interest information; updated title page with a note that this version of the manuscript has not been peer reviewed.

## References

1. Munoz-Price, L. S. et al. Clinical epidemiology of the global expansion of *Klebsiella pneumoniae* carbapenemases. Lancet Infect Dis 13, 785–796 (2013).

2. Kuehn, B. M. ‘Nightmare’ bacteria on the rise in US hospitals, long-term care facilities. JAMA 309, 1573–1574 (2013).

3. Murray, C. J. et al. Global burden of bacterial antimicrobial resistance in 2019: a systematic analysis. The Lancet 10.1016/S0140-6736(21)02724-0 (2022) doi:10.1016/S0140-6736(21)02724-0.

4. Wyres, K. L., Lam, M. M. C. & Holt, K. E. Population genomics of *Klebsiella pneumoniae*. Nat Rev Microbiol 18, 344–359 (2020).

5. Podschun, R. & Ullmann, U. *Klebsiella* spp. as Nosocomial Pathogens: Epidemiology, Taxonomy, Typing Methods, and Pathogenicity Factors. Clin Microbiol Rev 11, 589–603 (1998).

6. Martin, R. M. & Bachman, M. A. Colonization, Infection, and the Accessory Genome of *Klebsiella pneumoniae*. Frontiers in Cellular and Infection Microbiology 8, 4 (2018).

7. Martin, R. M. et al. Molecular Epidemiology of Colonizing and Infecting Isolates of *Klebsiella pneumoniae*. mSphere 1, e00261–16 (2016).

8. Gorrie, C. L. et al. Gastrointestinal Carriage Is a Major Reservoir of *Klebsiella pneumoniae* Infection in Intensive Care Patients. Clinical Infectious Diseases 65, 208–215 (2017).

9. Chung, D. R. et al. Fecal carriage of serotype K1 *Klebsiella pneumoniae* ST23 strains closely related to liver abscess isolates in Koreans living in Korea. Eur J Clin Microbiol Infect Dis 31, 481–486 (2012).

10. Lin, Y.-T. et al. Seroepidemiology of *Klebsiella pneumoniae* colonizing the intestinal tract of healthy Chinese and overseas Chinese adults in Asian countries. BMC Microbiol 12, 13 (2012).

11. Sun, Y. et al. Measurement of *Klebsiella* Intestinal Colonization Density To Assess Infection Risk. mSphere 6, e0050021 (2021).

12. Collingwood, A. et al. Epidemiological and Microbiome Associations Between *Klebsiella pneumoniae* and Vancomycin-Resistant *Enterococcus* Colonization in Intensive Care Unit Patients. Open Forum Infect Dis 7, ofaa012 (2020).

13. Vornhagen, J. et al. Combined comparative genomics and clinical modeling reveals plasmid-encoded genes are independently associated with *Klebsiella* infection. Nat Commun 13, 4459 (2022).

14. Diancourt, L., Passet, V., Verhoef, J., Grimont, P. A. D. & Brisse, S. Multilocus sequence typing of *Klebsiella pneumoniae* nosocomial isolates. J Clin Microbiol 43, 4178–4182 (2005).

15. Pan, Y.-J. et al. Capsular Types of *Klebsiella pneumoniae* Revisited by *wzc* Sequencing. PLOS ONE 8, e80670 (2013).

16. Brisse, S., et al. *wzi* Gene sequencing, a rapid method for determination of capsular type for *Klebsiella* strains. J Clin Microbiol 51, 4073–4078 (2013).

17. Lam, M. M. C., Wick, R. R., Judd, L. M., Holt, K. E. & Wyres, K. L. Kaptive 2.0: updated capsule and lipopolysaccharide locus typing for the *Klebsiella pneumoniae* species complex. Microb Genom 8, 000800 (2022).

18. Arlet, G. et al. Molecular epidemiology of *Klebsiella pneumoniae* strains that produce SHV-4 beta-lactamase and which were isolated in 14 French hospitals. J Clin Microbiol 32, 2553–2558 (1994).

19. Jolley, K. A. & Maiden, M. C. J. BIGSdb: Scalable analysis of bacterial genome variation at the population level. BMC Bioinformatics 11, 595 (2010).

20. Rao, K. et al. Risk Factors for *Klebsiella* Infections among Hospitalized Patients with Preexisting Colonization. mSphere 6, e0013221 (2021).

21. Rahn, A., Drummelsmith, J. & Whitfield, C. Conserved organization in the *cps* gene clusters for expression of Escherichia *coli* group 1 K antigens: relationship to the colanic acid biosynthesis locus and the cps genes from *Klebsiella pneumoniae*. J Bacteriol 181, 2307–2313 (1999).

22. Rahn, A., Beis, K., Naismith, J. H. & Whitfield, C. A novel outer membrane protein, Wzi, is involved in surface assembly of the *Escherichia coli* K30 group 1 capsule. J Bacteriol 185, 5882–5890 (2003).

23. Vornhagen, J., Rao, K. & Bachman, M. A. Gut community structure as a risk factor for infection in *Klebsiella pneumoniae*-colonized patients. mSystems 9, e0078624 (2024).

24. Diago-Navarro, E. et al. Carbapenem-resistant *Klebsiella pneumoniae* exhibit variability in capsular polysaccharide and capsule associated virulence traits. J Infect Dis 210, 803–813 (2014).

25. Wang, Y., Zhao, Y., Bollas, A., Wang, Y. & Au, K. F. Nanopore sequencing technology, bioinformatics and applications. Nat Biotechnol 39, 1348–1365 (2021).

26. Seekatz, A. M. et al. Presence of multiple *Clostridium difficile* strains at primary infection is associated with development of recurrent disease. Anaerobe 53, 74–81 (2018).

27. Caballero, S. et al. Distinct but Spatially Overlapping Intestinal Niches for Vancomycin-Resistant *Enterococcus faecium* and Carbapenem-Resistant *Klebsiella pneumoniae*. PLoS Pathog 11, e1005132 (2015).

28. Warren, R. M. et al. Patients with active tuberculosis often have different strains in the same sputum specimen. Am J Respir Crit Care Med 169, 610–614 (2004).

29. Nathavitharana, R. R. et al. Polyclonal Pulmonary Tuberculosis Infections and Risk for Multidrug Resistance, Lima, Peru. Emerg Infect Dis 23, 1887–1890 (2017).

30. Osbelt, L., et al. *Klebsiella oxytoca* causes colonization resistance against multidrug-resistant *K. pneumoniae* in the gut via cooperative carbohydrate competition. Cell Host Microbe 29, 1663–1679.e7 (2021).

31. Vezina, B. et al. A metabolic atlas of the Klebsiella pneumoniae species complex reveals lineage-specific metabolism that supports co-existence of diverse lineages. 2024.07.24.605038 Preprint at 10.1101/2024.07.24.605038 (2025).

32. Calderon-Gonzalez, R. et al. Modelling the Gastrointestinal Carriage of *Klebsiella pneumoniae* Infections. mBio 14, e03121–22 (2023).

33. Bray, A. S., et al. *Klebsiella pneumoniae* employs a type VI secretion system to overcome microbiota-mediated colonization resistance. Nat Commun 16, 940 (2025).

34. Vornhagen, J. et al. A plasmid locus associated with *Klebsiella* clinical infections encodes a microbiome-dependent gut fitness factor. PLOS Pathogens 17, e1009537 (2021).

35. Nanayakkara, B. S., O’Brien, C. L. & Gordon, D. M. Diversity and distribution of *Klebsiella* capsules in *Escherichia coli*. Environ Microbiol Rep 11, 107–117 (2019).

36. De Coster, W., D’Hert, S., Schultz, D. T., Cruts, M. & Van Broeckhoven, C. NanoPack: visualizing and processing long-read sequencing data. Bioinformatics 34, 2666–2669 (2018).

37. Li, H. Minimap2: pairwise alignment for nucleotide sequences. Bioinformatics 34, 3094–3100 (2018).

38. Bolger, A. M., Lohse, M. & Usadel, B. Trimmomatic: a flexible trimmer for Illumina sequence data. Bioinformatics 30, 2114–2120 (2014).

39. Olm, M. R. et al. inStrain profiles population microdiversity from metagenomic data and sensitively detects shared microbial strains. Nat Biotechnol 39, 727–736 (2021).

40. Li, H. et al. The Sequence Alignment/Map format and SAMtools. Bioinformatics 25, 2078–2079 (2009).

41. Schloss, P. D. et al. Introducing mothur: open-source, platform-independent, community-supported software for describing and comparing microbial communities. Appl Environ Microbiol 75, 7537–7541 (2009).

42. Quast, C. et al. The SILVA ribosomal RNA gene database project: improved data processing and web-based tools. Nucleic Acids Res 41, D590–596 (2013).

43. Oksanen, J. et al. vegan: Community Ecology Package. (2025).

44. Bursac, Z., Gauss, C. H., Williams, D. K. & Hosmer, D. W. Purposeful selection of variables in logistic regression. Source Code Biol Med 3, 17 (2008).

